# Development of Potent and Selective CK1α Molecular Glue Degraders

**DOI:** 10.1101/2024.10.01.616159

**Authors:** Qixiang Geng, Zixuan Jiang, Woong Sub Byun, Katherine A. Donovan, Zhe Zhuang, Fen Jiang, Hannah M. Jones, Hlib Razumkov, Michelle T. Tang, Roman C. Sarott, Eric S. Fischer, Steven M. Corsello, Stephen M. Hinshaw, Nathanael S. Gray

**Affiliations:** Department of Chemical and Systems Biology, Stanford Cancer Institute, School of Medicine, Stanford University, Stanford, CA USA; Sarafan ChEM-H, Stanford, CA USA; Department of Chemistry, Stanford School of Humanities and Sciences and Department of Chemical and Systems Biology, ChEM-H, Stanford School of Medicine, Stanford University, Stanford, CA USA; Department of Cancer Biology, Dana-Farber Cancer Institute, Boston, MA USA and Department of Biological Chemistry and Molecular Pharmacology Harvard Medical School, Boston, MA USA

## Abstract

Molecular glue degraders (MGDs) are small molecules that facilitate proximity between a target protein and an E3 ubiquitin ligase thereby inducing target protein degradation. Glutarimide-containing compounds are MGDs that bind to cereblon (CRBN) and recruit neosubstrates. Through explorative synthesis of a glutarimide-based library, we discovered a series of molecules that induce casein kinase 1 alpha (CK1α) degradation. By scaffold hopping and rational modification of the chemical scaffold, we identified an imidazo[1,2-a]pyrimidine compound that induces potent and selective CK1α degradation. A structure-activity relationship study of the lead compound, QXG-6442, identified the structural features that contribute to degradation potency and selectivity compared to other frequently observed neosubstrates. The glutarimide library screening and structure activity relationship medicinal chemistry approach we employed is generally useful for developing new molecular glue degraders towards new targets of interest.

## Introduction

Molecular glue degraders (MGDs) constitute a specialized class of small molecule that induce proximity between a target protein of interest and a component of the ubiquitin ligase machinery.[1–3] MGDs have two major advantages relative to bivalent proteolysis targeting chimera (PROTAC) degraders. The first is that they are typically smaller and to possess more ‘drug-like’ properties[2] which speeds medicinal chemistry optimization towards clinical candidates. The second is that they do not require a traditional ligand binding pocket on the protein of interest; many MGD targets were formerly considered to be very challenging to engage.[4] The major obstacle in MGD discovery is that the lack of systematic approaches to their discovery and optimization towards a given target of interest. [5]

We explore an established drug discovery approach: diversification and generalization of a privileged scaffold.[6,7] To implement this, we built combinatorial libraries of glutarimide-containing compounds.[8–10] This moiety is privileged in its ability to bind to a tri-tryptophan containing pocket of the CRL4^CRBN^ ubiquitin ligase. Multiple serendipitously discovered glutarimide derivates, including the immunomodulatory drugs (IMiDs) such as thalidomide, lenalidomide, and pomalidomide, demonstrate the potential of this compound class. These classic examples are currently used for the treatment of multiple myeloma.[11,12] The aforementioned IMiDs, derive their shared pharmacology from the ability to induce ubiquitin-mediated degradation of the Ikaros family zinc finger transcription factors (IKZF1 and IKZF3) through recruitment of the CRL4^CRBN^ E3 ligase.[13–15] Despite their structural similarity, IMiDs can produce completely distinct protein degradation profiles beyond IKZF1/3. For example, among these three FDA-approved IMiDs, only lenalidomide induces the degradation of casein kinase 1 alpha (CK1α),[16] highlighting how subtle chemical modifications can influence neosubstrate recruitment and degradation (Figure 1A).[17]

**Figure 1.**
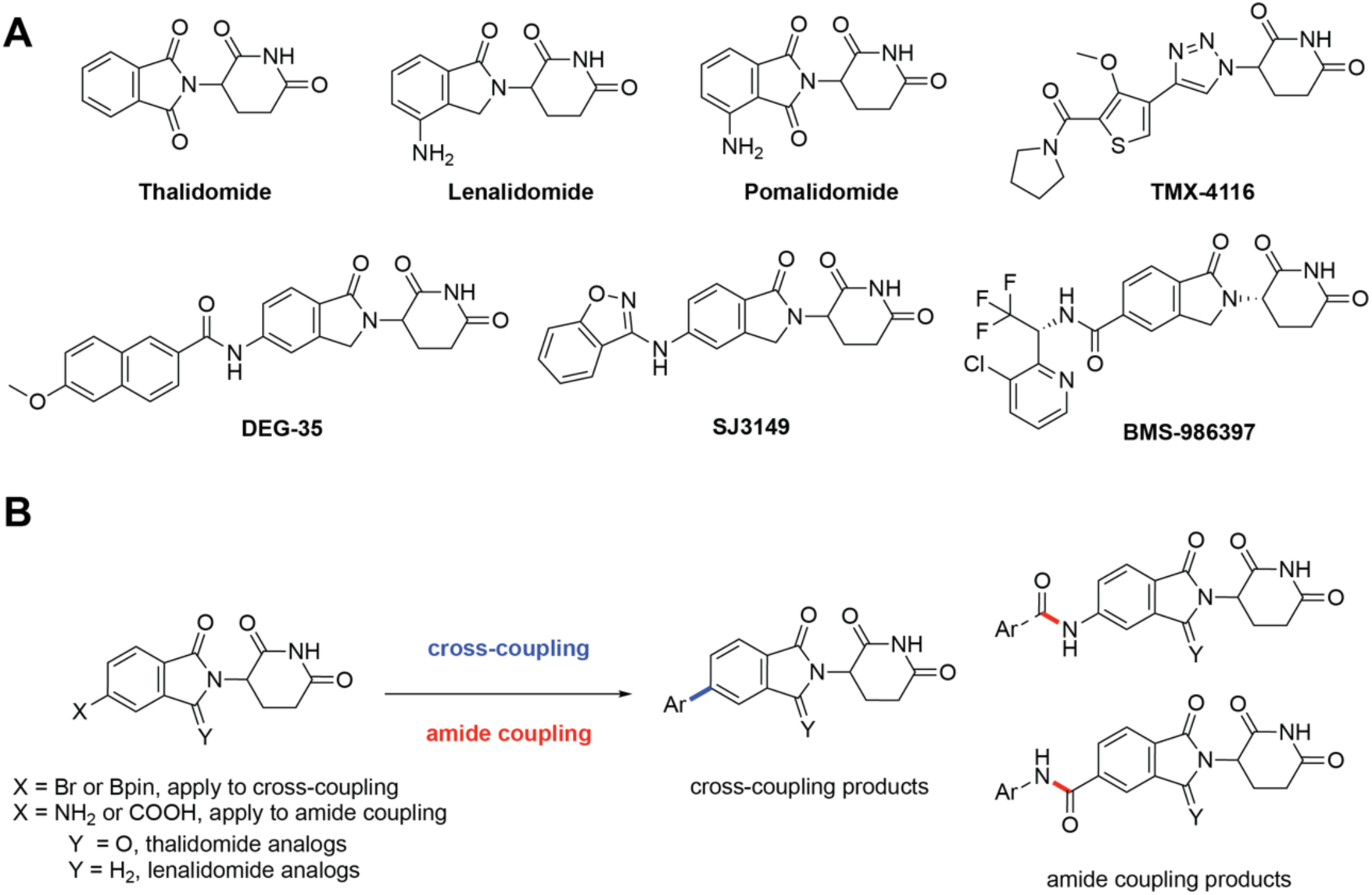
Chemical structures of IMiDs and library design. (A) Chemical structure of representative IMiDs known to degrade CK1α. (B) Library design based on two chemical strategies; Ar = Aromatic group.

CK1α is a potential therapeutic target in acute myeloid leukemia (AML), a frequently fatal bone marrow stem cell cancer with high mortality. AML is the most common malignant myeloid disorder in adults.[18,19] AML cell lines are sensitive to p53 activation,[20] which can be triggered by silencing of CK1α via inhibition, genetic knockdown, or degradation.[21,22] While lenalidomide induces weak and non-selective CK1α degradation in AML cells, more selective and potent compounds are needed to fully evaluate the therapeutic potential of degrading or inhibiting this target.[23] Several selective degraders of CK1α have been reported. Our laboratory identified **TMX-4116**, a selective CK1α MGD, during the development of PDE6D degraders (Figure 1A).[10] More recently, Woo and colleagues reported a dual degrader of IKZF2 and CK1α, **DEG-35**, that exhibits single digit nanomolar level cytotoxicity against AML cell lines.[24,25] Rankovic and colleagues reported the development of **SJ3149**, which degrades CK1α potently *in vitro* and in mice, has also been reported. In this study, a crystal structure of the CK1α+**SJ3149**+CRBN+DDB1 quaternary complex enabled visualization of the key interactions required for molecular recognition[26](Figure 1A). Finally, a CK1α MGD from BMS, CC-91633 (**BMS-986397**), entered Phase 1 clinical trials (NCT04951778). Given the promising preclinical study results, defining the structural characteristics of glutarimide-based MGDs responsible for selectivity and potency against CK1α is highly important for drug discovery.

Development of selective and potent CRBN-dependent MGDs follows an established path. We previously reported GSPT1 MGDs that were identified from a focused combinatorial library enabled by nucleophilic aromatic substitution (SNAr).[8] The construction of an IMiD library through click chemistry and the Groebke-Blackburn-Bienaymé (GBB) reaction enabled the discovery of selective MGDs for phosphodiesterase 6D (PDE6D)[10] and WEE1[9], respectively. Herein, we use two of the most frequently used reactions in medicinal chemistry,[27] namely metal-mediated cross-coupling and amide coupling, for the efficient construction of an IMiD library with over 100 compounds. We installed heterocyclic moieties[28] on either the phthalimide skeleton of thalidomide or the isoindolinone skeleton of lenalidomide analogs (Figure 1 A, B). Screening of the cross-coupling-derived IMiD library led to the discovery of **QXG-0629**, a potent degrader of CK1α. GSPT1, WEE1, and IKZF2 were additional targets. Further derivatization of **QXG-0629** resulted in the discovery of **QXG-0632** and **QXG-6442**, which display potent and selective CK1α degradation activity. These findings highlight the potential of our innovative approach to unlock new therapeutic avenues in targeting CK1α and advancing the fight against acute myeloid leukemia.

## Results and Discussion

### Library synthesis and discovery of selective CK1α degraders

We synthesized two libraries of glutarimide analogs: the first library was accessed by palladium-catalyzed cross coupling of substituted thalidomide derivatives, while the second was generated using amide bond formation reactions (Figure 1B). As part of a larger discovery campaign to identify new MGDs, we performed quantitative mass spectrometry (MS)-based proteomics in MOLT-4 cells using these libraries. From this screening, we first identified **QXG-0629**, which has an isoquinoline attached from its 3-position to the isoindolinone moiety of lenalidomide analogs via a C-C bond. This compound degraded CK1α, GSPT1, WEE1 and IKZF2 after 5 h treatment at a concentration of 1 μM in MOLT-4 cells (Figure 2A, B). Validation by Western blot analysis confirmed these findings (Figure 2D).

**Figure 2.**
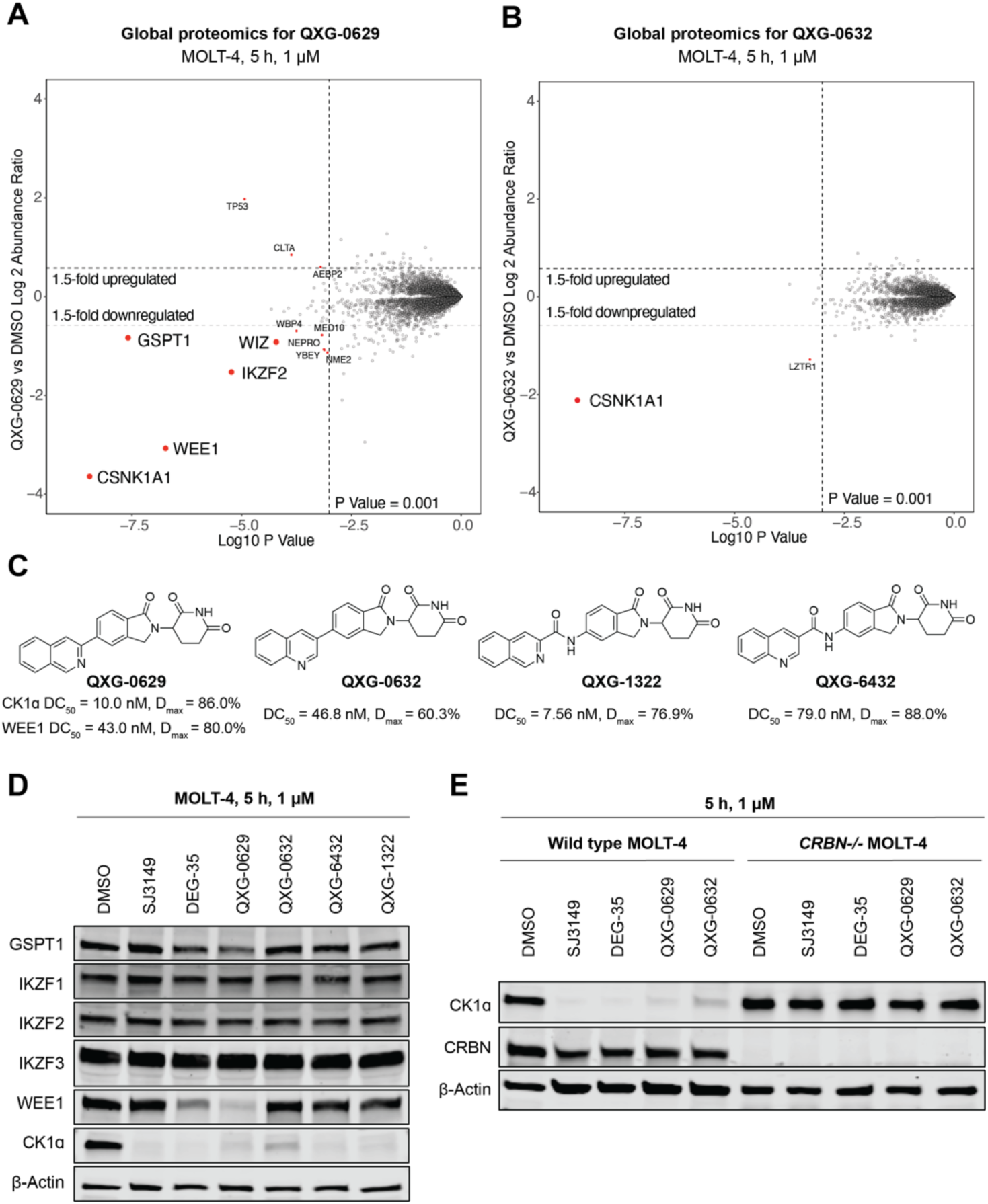
Discovery of compounds selectively degrade CK1α in a CRBN-dependent manner. (A) Global proteomics analysis of **QXG-0629** at 1 μM in MOLT-4 cells after 5 h treatment. (B) Global proteomics analysis of **QXG-0632** at 1 μM in MOLT-4 cells after 5 h treatment. (C) Chemical structures of **QXG-0629**, **QXG-0632**, **QXG-6432** and **QXG-1322**. (D) Western blot analysis showing degradation effects of compounds on known CRBN neosubstrates at 1 μM in MOLT-4 cells after 5 h treatment. (E) Western blot analysis of CK1α in wildtype and *CRBN-/-* MOLT-4 cells treated with 1 μM of **QXG-0629** and **QXG-0632** for 5 h.

To effectively test CK1α degrader activity in higher-throughput, we developed a CK1α-HiBiT assay by tagging the endogenous protein to quantitatively measure protein abundance. In parallel, we employed a WEE1-HiBiT assay as a counter-screen to evaluate degradation selectivity. HiBiT assays leverage a split luciferase system to monitor protein abundance and are now routinely used in the targeted protein degradation field for target-centric screening.[29] In these assays, **QXG-0629** demonstrated a slightly lower half-maximal degradation (DC_50_) value of 10.0 nM for CK1α compared to the DC_50_ of 43.0 nM for WEE1, yet both achieved a similar maximum target fraction degraded (D_max_) (Figure 2C). To improve degradation selectivity through rational modification, we initiated a medicinal chemistry campaign by modifying the heteroaromatic ring and varying the distance between the gluing moiety and the CRBN-binding moiety. Both types of chemical modifications resulted in the discovery of selective and potent CK1α degraders bearing only minor structurally changes: **QXG-0632** contains a quinoline instead of isoquinoline while **QXG-6432** and **QXG-1322** contain amide insertions (Figure 2C). **QXG-0632** induced HiBiT-CK1α degradation with higher DC_50_ values (46.8 nM) than **QXG-0629**, with D_max_ of 60.3%, but no HiBiT-WEE1 degradation was observed with **QXG-0632** treatment at concentration ranging from 4.6 nM to 10 µM. Similar selectivity was observed with **QXG-1322** and **QXG-6432**, both of which exhibited improved CK1α degradation potency, with DC_50_ values of 7.5 nM (D_max_ = 76.9%) and 79.0 nM (D_max_ = 88.0%), respectively. Western blot validation confirmed that all of these three new analogs selectively depleted CK1α in MOLT-4 cells at 1 µM within 5 h without degrading WEE1, GSPT1, or IKZF1/3 (Figure 2D). **QXG-0632**, which contains the quinoline isomer of **QXG-0629**, exhibits proteome-wide selectivity for degradation of CK1α (Figure 2B). All degraders induce CK1α degradation in a CRBN dependent manner, as demonstrated by the rescue of CK1α degradation with CRBN-knockout cells (Figure 2E).

Given that the introduction of an amide linker improved selectivity and potency, we focused our next SAR efforts on synthesizing additional analogs incorporating this modification. We discovered that introducing an additional nitrogen into the quinoline fragment to afford the corresponding 2-quinoxaline enhanced CK1α degradation potency to the single digit nM range (**QXG-1323**, DC_50_ = 9 nM), while only slightly reducing the observed D_max_ to 80%. Further contraction of the heterocycle resulted in **QXG-6442**, featuring a 2-imidazo[1,2-*a*]pyridine moiety, which exhibited a DC_50_ value of 5.7 nM and an improved D_max_ of 90%. Notably, **QXG-6442** exhibited impressive proteome-wide selectivity for CK1α degradation in MOLT-4 cells at a concentration of 1 μM (Figure 3A).

**Figure 3.**
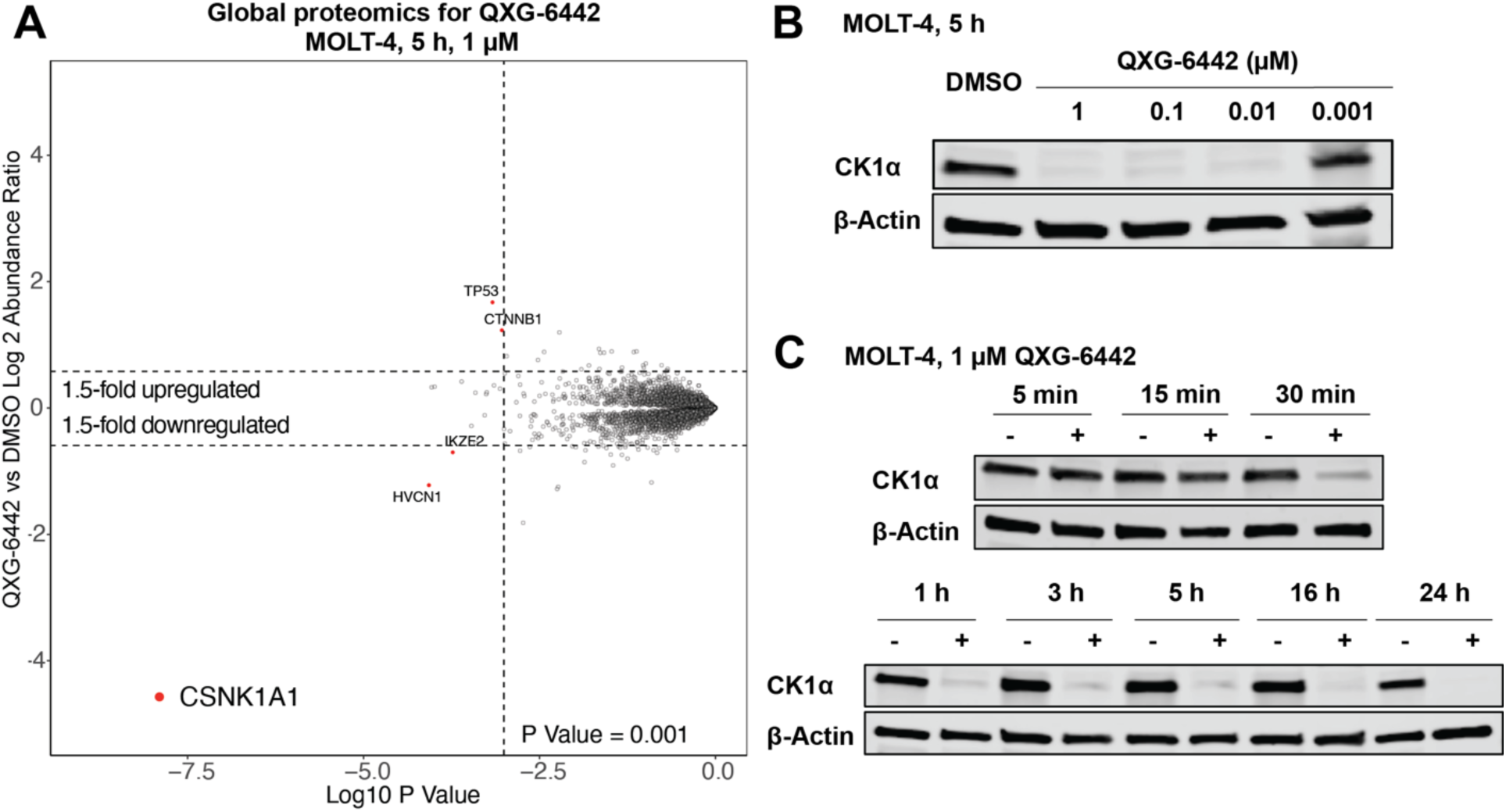
Characterization of the proteome-wide degradation selectivity of **QXG-6442.** (A) Global proteomics analysis of **QXG-6442** at 1 μM in MOLT-4 cells after 5 h treatment. (B) Western blot analysis of CK1α in MOLT-4 cells dosed with **QXG-6442** for 5 h. (C) Timepoint-dependent Western blot analysis of CK1α treated with 1 μM of **QXG-6442**.

Based on these results, we initiated a SAR study on the glutarimide moiety, the pharmacophore responsible for binding to CRBN. We first synthesized a methylated compound, **QXG-6142**, a modification that is well-known to disrupt a key hydrogen bond to CRBN, as a potential negative control compound (Table 1). As expected, no degradation was observed in the HiBiT assay, in agreement with the Western blot results in Figure 2E. Next, we synthesized **QXG-6471** and **QXG-6477** (Table 1), which have a phthalimide core instead of the isoindolinone motif present in **QXG-6442** and **QXG-6432**. This structural change significantly reduced CK1α degradation activity, indicating that the isoindolinone is required for the observed potency. Moreover, the analogs with an additional methylene group derived from **QXG-6432** and **QXG-6442** (**QXG-6431** and **QXG-6441** in Table 1) lost the ability to degrade CK1α, demonstrating the importance of the distance between the CRBN-binding subunit and the heterocyclic moiety. Additionally, the 4- or 6-substituted IMiDs (**QXG-6433** and **QXG-6113**) all failed to degrade CK1α, suggesting that the substitution position on the isoindolinone motif influences the potency (Figure 3A). Nonetheless, **QXG-0663**, with 4-substitution but also possessing an additional methylene group can still degrade CK1α. This suggests that increased flexibility of the gluing moiety may partially compensates for the substitution effect.

**Table 1.**
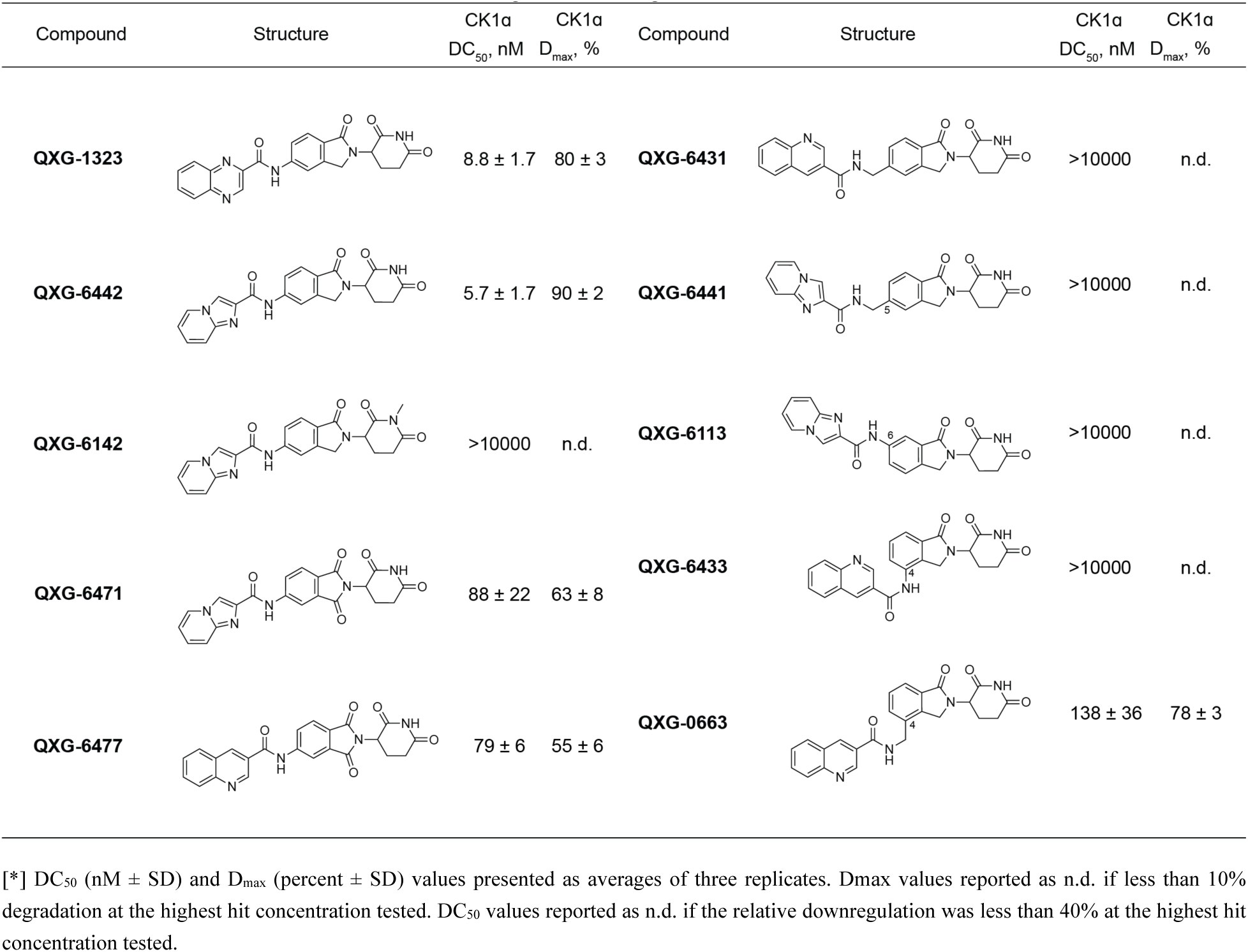
Structures of amide-based derivates derived from **QXG-1322** and **QXG-6432**.*

| Compound | Structure | CK1a<br>DC<sub>50</sub>, nM | CK1a<br>D<sub>max</sub>, % | Compound | Structure | CK1a<br>DC<sub>50</sub>, nM | CK1a<br>D<sub>max</sub>, % |
| --- | --- | --- | --- | --- | --- | --- | --- |
| **QXG-1323** |  | 8.8 ± 1.7 | 80 ± 3 | **QXG-6431** |  | >10000 | n.d. |
| **QXG-6442** |  | 5.7 ± 1.7 | 90 ± 2 | **QXG-6441** |  | >10000 | n.d. |
| **QXG-6142** |  | >10000 | n.d. | **QXG-6113** |  | >10000 | n.d. |
| **QXG-6471** |  | 88 ± 22 | 63 ± 8 | **QXG-6433** |  | >10000 | n.d. |
| **QXG-6477** |  | 79 ± 6 | 55 ± 6 | **QXG-0663** |  | 138 ± 36 | 78 ± 3 |

[\*] DC<sub>50</sub> (nM ± SD) and D<sub>max</sub> (percent ± SD) values presented as averages of three replicates. Dmax values reported as n.d. if less than 10% degradation at the highest hit concentration tested. DC<sub>50</sub> values reported as n.d. if the relative downregulation was less than 40% at the highest hit concentration tested.

We further analyzed the ability of **QXG-0632** and **QXG-6442** to degrade CK1α in MOLT-4 cells (Figure 3B, Figure S1). A time-course analysis demonstrated that **QXG-6442** and **QXG-0632** can degrade CK1α in less than 3 h at a concentration of 1 μM. **QXG-6442** also exhibited improved CK1α degradation within 30 min (Figure 3C, Figure S2).

To rationalize the observed structure activity relationships we performed molecular docking analysis of **QXG-0632** and **QXG-6442**, informed by the high resolution ternary complex structure of **SJ3149** with CK1α and CRBN-DDB1 (Figure 4).[26] The docking model indicates the left-side binary ring system of both compounds inserted into a hydrophobic groove of CK1α and forms hydrophobic interactions with VAL20 and ILE35. It also predicts that the improved recognition of CK1α by **QXG-0632** may result from the isoquinoline moiety that forms a cation-π interaction between the benzo-section of the quinoline motif and LYS18 of CK1α. The further optimized compound **QXG-6442** can potentially form a hydrogen bond between the 1-nitrogen of the imidazo[1,2-*a*]pyridine motif and the same LYS18. Similarly, cation-π interaction between the pyridinyl-section of the quinoline motif and the HIS353 contributes to the improved affinity to CRBN. While the same histidine formed an additional hydrogen bond with the carbonyl group of **QXG-6442**, potentially explaining its ability to induce stronger degradation than **QXG-0632**. This hydrogen bond is also observed for **SJ3149** in the experimentally determined quaternary complex (Ref PDB: 8G66). The enhanced interactions of **QXG-6442** with CK1α and CRBN likely contribute to the improved binding affinity, and high potency observed in ternary complex formation and degradation. The docking models also indicated that the left-side binary ring system has potential for further modification to improve the properties of the compounds.

**Figure 4.**
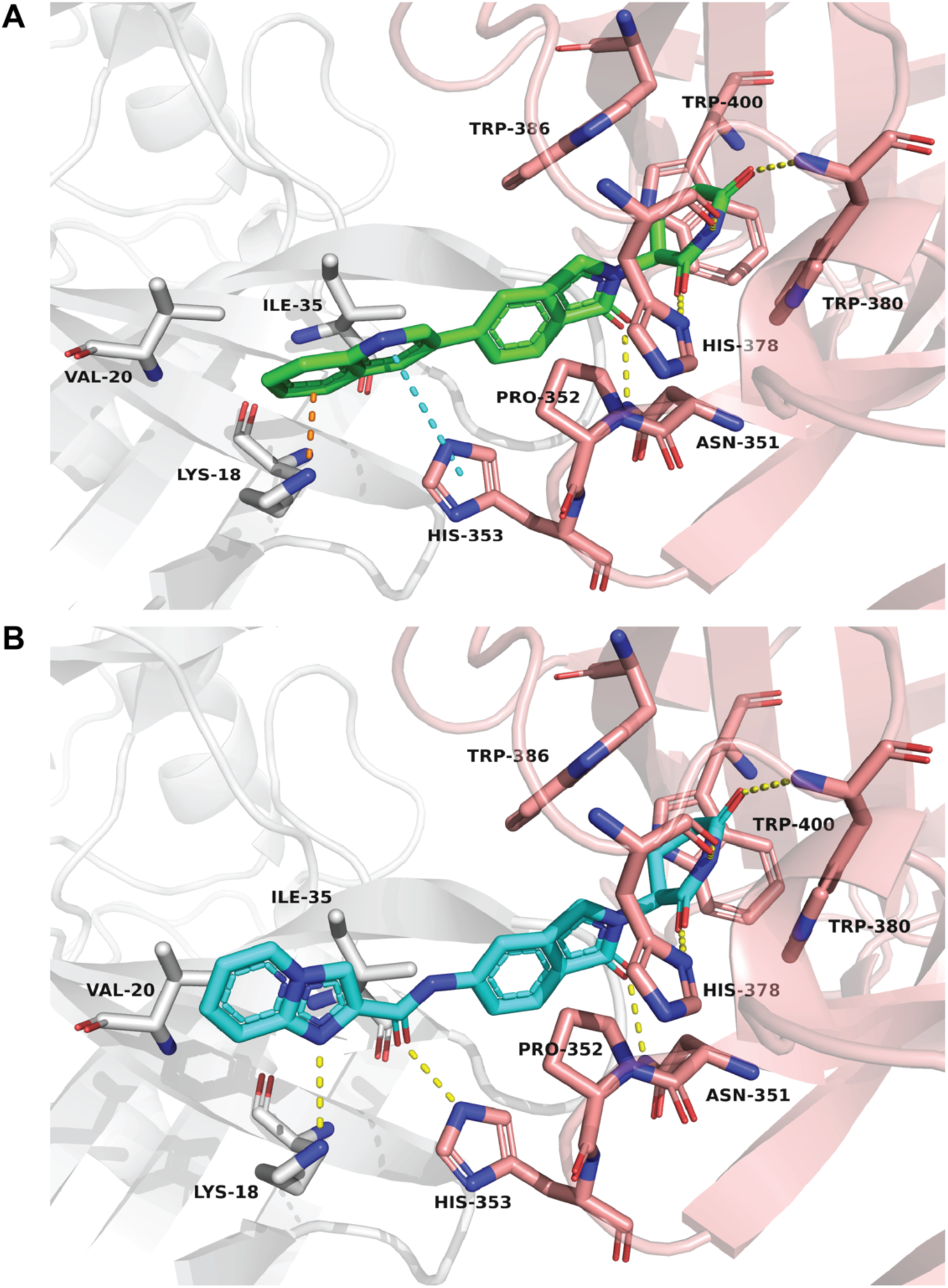
Docking model of CRBN-**QXG-0632**-CK1α and CRBN-**QXG-6442**-CK1α ternary complex. Compounds and the residues that involved in the binding are shown in sticks. CK1α, CRBN, **QXG-0632** and **QXG-6442** are colored in grey, pink, green and blue, respectively.

### SAR study of heterocyclic gluing moiety

Having identified the selective CK1α degrader **QXG-6442**, we sought to further explore the chemical space of the heterocyclic gluing moiety. First, we changed the carbonyl group attachment of the imidazo[1,2-*a*]pyridine motif from the 2-position to the 3-position, resulting in **QXG-131-5**. This adjustment resulted in a greater than 10-fold decrease in degradation potency, indicating the importance of the 2-position connection of the imidazo[1,2-*a*]pyridine skeleton. Next, we rearranged the two nitrogens on the heterocycle, leading to the synthesis of **QXG-133-3** with a pyrazole and **QXG-133-6** with a 1H-benzimidazole. These modifications retained degradation activity, suggesting that 1-nitrogen of the imidazo[1,2-*a*]pyridine, which is predicted by the docking model to provide a hydrogen bond acceptor with LYS18 of CK1α. As such, the 1-nitrogen may have contributed predominating to the observed activity. To confirm the importance of the 1-nitrogen we replaced it with a carbon to afford the indolizine analog **QXG-133-2**. As expected, the loss of one more hydrogen bond resulted in more than a 10-fold increase in DC_50_ and decreased D_max_ values. On the other hand, since adding an additional nitrogen leads to a potency improvement from **QXG-0632** to **QXG-1323** and **QXG-6442** (Table 1), addition of nitrogen in **QXG-6442** and **QXG-133-3** resulted in **QXG-133-4** and **QXG-133-5**. (Table 2) Unfortunately, both modifications also resulted in a greater than 10-fold decrease in degradation potency, potentially because the increased electron density weakens the ability of 1-nitrogen as a hydrogen bond acceptor as suggested by docking.

**Table 2.** Structure-activity relationship of heterocyclic gluing moiety.*

| Compound | Structure | CK1a<br>DC<sub>50</sub>, nM | CK1a<br>D<sub>max</sub>, % |
| --- | --- | --- | --- |
| **QXG-131-5** |  | 82 ± 13 | 83 ± 6 |
| **QXG-133-3** |  | 4.7 ± 0.5 | 85 ± 4 |
| **QXG-133-6** |  | 11.1 ± 0.7 | 84 ± 1 |
| **QXG-133-2** |  | 87 ± 23 | 79 ± 8 |
| **QXG-133-4** |  | 61 ± 10 | 83 ± 1 |
| **QXG-133-5** |  | 61 ± 4 | 85 ± 4 |

[\*] DC<sub>50</sub> (nM ± SD) and D<sub>max</sub> (percent ± SD) values presented as averages of three replicates. Dmax values reported as n.d. if less than 10% degradation at the highest hit concentration tested. DC<sub>50</sub> values reported as n.d. if the relative downregulation was less than 40% at the highest hit concentration tested.

Focusing on **QXG-6442**, we further explored the substituents on the imidazo[1,2-*a*]pyridine motif. **QXG-131-6** with 3-methyl, **QXG-131-7** with 5-methyl, and **QXG-131-3** with 8-methoxyl group maintained similar degradation potency, indicating steric tolerance around this moiety (Table 3). We also investigated inductive effect of either electron donating or withdrawing groups on activity, by synthesizing a series of methoxy- or halogen-containing compounds. We found that all these substituents are well tolerated. Comparing the potency of **QXG-131-2**, **QXG-131-1** and **QXG-131-3**, with methoxy group at the 6-, 7- and 8-positions respectively, to **QXG-6442**, we found that the methoxy group modification can improve the activity. Specifically, the 6-substitution resulted in the lowest DC_50_ and the highest D_max_, indicating a possible hydrophobic interaction between the 6-methoxy group and CK1α. This modification is also structurally consistent with **DEG-35**. However, we found that using the imidazo[1,2-*a*]pyridine scaffold can further enhance activity. Replacing the methoxy- with a chloro-group (**QXG-131-8**) retained strong potency, but a fluorine substitution led to similar activity to the lead compound. Among fluorine substitutions, **QXG-131-10** with 7-fluorine resulted in weaker degradation than the corresponding 6-flouro substituted compound **QXG-131-9**. Replacing the methoxy group with a trifluoromethyl group at the 8-position in **QXG-131-4**, significantly improved the degradation potency compared with **QXG-131-3**. This indicates the possibility of a strong electron-withdrawing inductive effect on 1-nitrogen of imidazo[1,2-*a*]pyridine improves the hydrogen bonding to CK1α.

### Antiproliferative effects of CK1α degraders

We screened several potent CK1α degraders across a small panel of cancer cell lines to access their antiproliferative effects. Using **SJ3149** as a positive control, we tested **QXG-0632** and **QXG-6442** in 72 h Cell-Titer Glo (CTG) cell line proliferation assays spanning acute lymphoblastic leukemia (ALL) (MOLT-4, Jurkat), diffuse large B-cell lymphoma (DLBCL) (SU-DHL-5), acute promyelocytic leukemia (APL) (NB-4), and AML (NOMO-1, MOLM-13 and MOLM-14) (Figure S3). Our CK1α degraders exhibited promising antiproliferative effects in MOLM-13 and MOLM-14 cells (Figure 5A, B), comparable to **DEG-35** and **SJ3149**, likely due to the higher sensitivity of AML cells to CK1α levels. We also treated the wild-type and CRBN-knockout MOLM-14 cells with the representative degraders **QXG-0632**, **QXG-6442**, **QXG-131-2**, **QXG-131-4** and **QXG-133-3**. We observed that all these compounds induced antiproliferative effects in a CRBN-dependent manner (Figure 5B). Western blot analysis confirmed CK1α degradation in MOLM-13 and MOLM-14 cells after 5 h treatment with 1 μM of the degraders (Figure 5C). To understand the antiproliferative SAR of **QXG-6442** analogs in Table 2 and Table 3, we tested all these compounds using the CTG assay in MOLM-13 and MOLM-14 cells (Table S1 and Table S2). Our results revealed that the SAR for antiproliferative effect shows a similar trend to that of degradation. The CTG assay EC_50_ and HiBiT DC_50_ of the CK1α degraders derived from the imidazole[1,2-*a*]pyridine sceffold (Table 3) were correlated in MOLM-13 and MOLM-14 cells, indicating that the antiproliferative effect is likely to be caused by CK1α degradation (Figure S5). We further assessed the proteome-wide selectivity of **QXG-6442** in MOLM-13 cells, the most sensitive cell line, using global proteomics analysis. CK1α was the only protein that decreased after 5 h treatment with 1 μM drug, indicating high CK1α selectivity (Figure S6).

**Figure 5.**
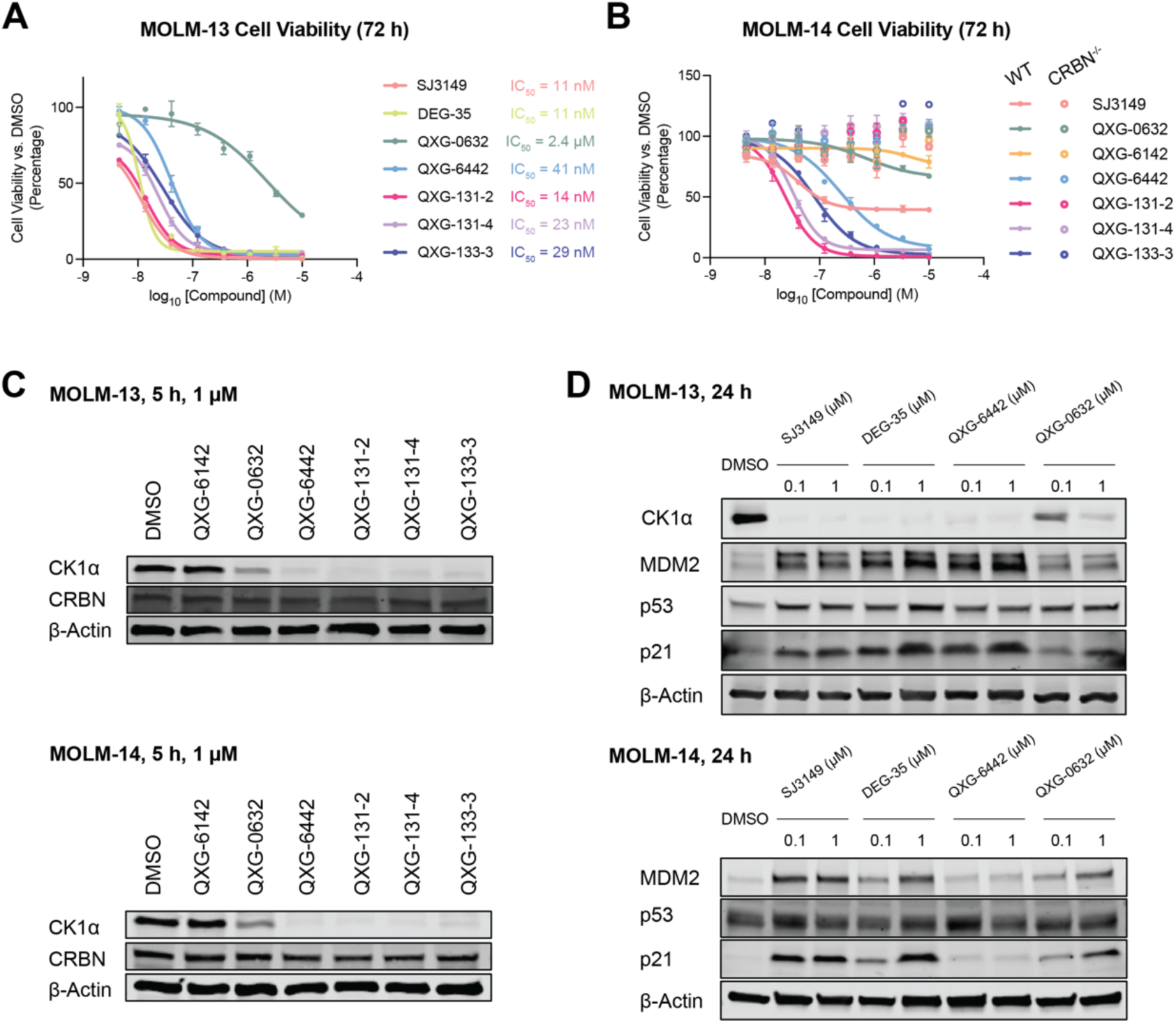
Antiproliferative activity of potent CK1α degraders in MOLM-13 and MOLM-14 cell lines and the downstream effects. (A) Antiproliferative screening of potent CK1α degraders in MOLM-13 cells: MOLM-13 cells were incubated with compounds, DMSO as a negative control and **SJ3149** as a positive control at the indicate range of concentrations (n = 3). After 72 h, cell proliferation was evaluated using the Cell-Titer Glo (CTG) assay. (B) Antiproliferative screening of potent CK1α degraders in MOLM-14 wild type and *CRBN -/-* MOLM-14 cells. Cells were incubated with a compound, DMSO as a negative control, **SJ3149** as a positive control at the indicate range of concentrations (n = 3). After 72 h, cell proliferation was evaluated by the CTG assay. (C) Western blot analysis of CK1α degradation in MOLM-13 and MOLM-14 cells treated for 5 h with 1 μM of CK1α degraders. (D) Western blot analysis of MDM2, p53, and p21 in MOLM-13 and MOLM-14 cells treated for 5 h with either 0.1 or 1 μM of CK1α degraders.

To explore the mechanism of the antiproliferative effects induced by degraders and evaluate the sensitivity across different cell lines, we selected MOLM-13 and MOLM-14 cells as models for studying downstream effects. By Western blotting, we measured the levels of p21, p53, and MDM2 levels after treating cells with 0.1 or 1 μM of **QXG-0632**, **QXG-6442**, and two control degraders, **DEG-35** and **SJ3149**, for 24 hours. As expected and consistent with a previous study,[26] increased levels of MDM2, p53, p21, and β-catenin were observed in both cell lines, while differing levels of p21 and MDM2 upregulation were observed across the compounds with different potency. (Figure 5D). These findings indicate that the antiproliferative effect is closely correlated with CK1α degradation, which plays a crucial role in regulation of β-catenin and p21 levels within the p53 pathway.[30]

### *In vitro* microsomal stability of selective CK1α degraders

We evaluated the in vitro metabolic stability and intrinsic reactivity of selected degraders to further evaluate their ADME properties. We chose the selective CK1α degraders **QXG-0632**, **QXG-6432**, **QXG-6442**, the activity-improved analog **QXG-131-2**, and the rearranged scaffold compound **QXG-133-3** as representative candidates. **QXG-0632** and **QXG-6442** exhibited good metabolic stability with half-life (T_1/2_) of more than 120 h with intrinsic clearance of less than 6 μL/min/mg in human liver microsomes. Additionally, both compounds also exhibited reasonable T_1/2_ of 65.6 min and 75.4 min and intrinsic clearance of 11 and 9 μL/min/mg, respectively in mouse liver (Table 4). Among the compounds developed in the SAR studies, only imidazo[1,2-*a*] featuring compound **QXG-6442**, maintained high stability and low clearance in both mouse and human liver microsomes compared to the initial lead compound **QXG-0632**. Addition of a methoxy modification in **QXG-131-2** led to a significant reduction in half-life in liver and increased intrinsic clearance. Thus, **QXG-6442** is a promising candidate for further experiments in rodents.

**Table 3.**
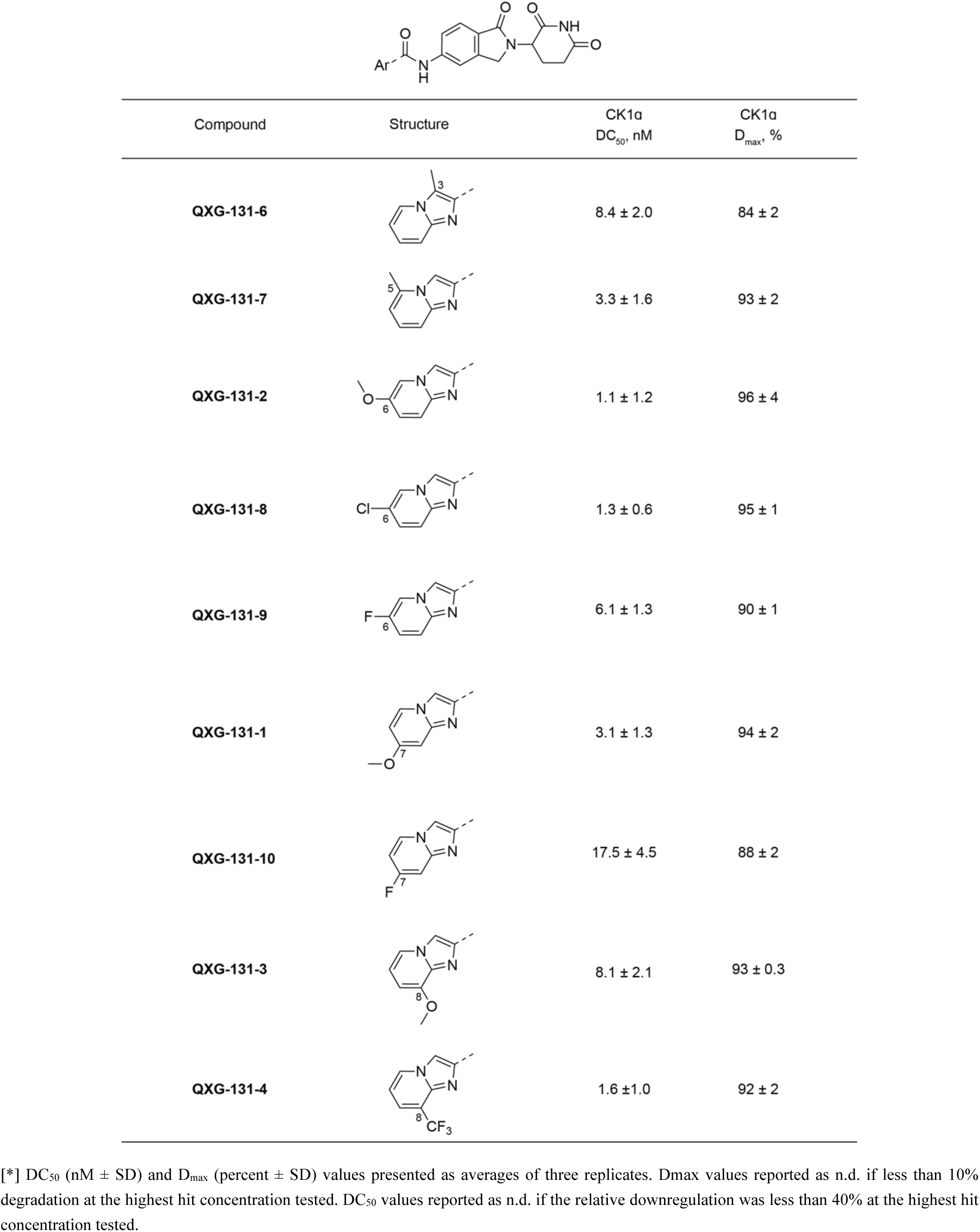
Structure-activity relationship of the substituents on the imidazo[1,2-a]pyridine moiety.*

| Compound | Structure | CK1a<br>DC<sub>50</sub>, nM | CK1a<br>D<sub>max</sub>, % |
| --- | --- | --- | --- |
| **QXG-131-6** |  | 8.4 ± 2.0 | 84 ± 2 |
| **QXG-131-7** |  | 3.3 ± 1.6 | 93 ± 2 |
| **QXG-131-2** |  | 1.1 ± 1.2 | 96 ± 4 |
| **QXG-131-8** |  | 1.3 ± 0.6 | 95 ± 1 |
| **QXG-131-9** |  | 6.1 ± 1.3 | 90 ± 1 |
| **QXG-131-1** |  | 3.1 ± 1.3 | 94 ± 2 |
| **QXG-131-10** |  | 17.5 ± 4.5 | 88 ± 2 |
| **QXG-131-3** |  | 8.1 ± 2.1 | 93 ± 0.3 |
| **QXG-131-4** |  | 1.6 ± 1.0 | 92 ± 2 |

[\*] DC<sub>50</sub> (nM ± SD) and D<sub>max</sub> (percent ± SD) values presented as averages of three replicates. Dmax values reported as n.d. if less than 10% degradation at the highest hit concentration tested. DC<sub>50</sub> values reported as n.d. if the relative downregulation was less than 40% at the highest hit concentration tested.

**Table 4.** Metabolic stability of selected degraders.

| Compound | Species (T<sub>1/2</sub> in minutes) |  | Species (Cl<sub>int</sub> in µ/min/mg) |  |
| --- | --- | --- | --- | --- |
|  | Human | Mouse | Human | Mouse |
| Sunitinib | 39.4 | 20.9 | 18 | 33 |
| **QXG-0632** | >120 | 65.6 | <6 | 11 |
| **QXG-6442** | >120 | 75.4 | <6 | 9 |
| **QXG-6432** | 23.4 | >120 | 30 | <6 |
| **QXG-131-2** | 33.2 | >120 | 21 | <6 |
| **QXG-133-3** | 65.4 | 6.3 | 11 | 111 |

## Conclusion

We developed selective and potent CK1α MGDs synthesized by modification of the isoindolinone core of lenalidomide using cross-coupling and amide-coupling reactions. Initial screening of over 100 IMiD analogs resulted in the hit CK1α degrader **QXG-0629**, which also degraded WEE1, GSPT1, and IKZF2. Through chemical modification of this hit compound, we developed several improved CK1α degraders, including **QXG-6442**. Further SAR exploration of the imidazo[1,2-*a*]pyridine scaffold in the lead compound **QXG-6442** indicated good tolerance towards substituents on the neosubstrate-contacting heterocycle, establishing a series of potent analogs with single-digit nanomolar activity. Consistent with the selective CK1α degraders developed by others,[24,26] our structural docking model predicted the key hydrophobic interactions and hydrogen bonds of **QXG-6442** with CK1α and CRBN, providing a clear rationale for the observed SAR. This also opens opportunities for further structural modifications in drug development.

The antiproliferative effect of the CK1α degraders varied across different cell lines. AML cell lines showed the greatest sensitivity, and we observed stabilization of p53 and induction of p53 regulated genes including MDM2 and p21. Additionally, these new CK1α degraders also exhibited good microsomal stability suggesting that they represent promising leads for further optimization. We anticipate that the tool compounds developed in this study will be useful as pharmacological probes of CK1α degradation dependent pharmacology.

## Methods

### Cell Lines

Jurkat (ATCC, #TIB-152) and MOLT-4 (ATCC, #CRL-1582) were maintained in RPMI-1640 (Gibco) supplemented with 10% heat-inactivated fetal bovine serum (FBS, Gibco) and 1% penicillin-streptomycin (Gibco). MOLM-13 (DSMZ, ACC 554) and MOLM-14 (DSMZ, ACC 777) were maintained in RPMI-1640 (Gibco) supplemented with 20% heat-inactivated fetal bovine serum (FBS, Gibco) and 1% penicillin-streptomycin (Gibco). All cell lines were cultured at 37 °C with 5% CO_2_ in a humidified atmosphere for no longer than 4 weeks for all experiments. Mycoplasma testing was performed monthly using the MycoAlert mycoplasma detection kit (Lonza, Basel, Switzerland).

### Cell Proliferation Assays (CellTiter-Glo)

Cells were seeded at a density of 1.0 × 10^3^ cells per well in 384-well opaque plates (Corning) containing 50 µL of growth medium. Drug treatments were applied immediately after seeding at the indicated concentrations. Cellular ATP levels, as an indicator of cell viability, were measured 72 h post-treatment using the CellTiter-Glo Luminescent Cell Viability Assay (Promega) as described in manufacturer’s manual. The half-maximal inhibitory concentration (IC₅₀) values were calculated from nonlinear regression analysis of dose-response curves generated in GraphPad Prism 10.2.2, with each data point representing an independent experimental replicate. IC₅₀ values are presented as the mean of the IC_50_ values from the singlet fits.

### HiBiT Assays

Introduction of a HiBiT coding sequence into the endogenous *Wee1/CK1α* locus in Jurkat cells was done via CRISPR-Cas9 genome editing as being described.^1^ HiBiT-tagged Jurkat cells (2.0 × 10^4^ cells per well) were seeded in 50 µL of growth medium in opaque white 384-well plates (Corning). Compounds were dispensed using a D300e digital dispenser (Tecan) from DMSO stock solutions to achieve the indicated final concentrations. The cells were incubated at 37 °C for 5 h. Endogenous protein levels were quantified using the Nano-Glo HiBiT Lytic Detection System (Promega) as described in manufacturer’s manual. All experiments were conducted in triplicate and repeated at least twice on separate days. DC₅₀ values were calculated through nonlinear regression analysis using GraphPad Prism 10.2.2, with each set of singlet measurements treated as independent replicates for curve fitting.

### Immunoblotting

Whole cell lysates for immunoblotting were prepared by pelleting cells from each cell line at 500 g for 5 min at 4 °C. The cell pellets were suspended in RIPA Lysis and Extraction Buffer (Thermo Fisher Scientific) supplemented with protease and phosphatase inhibitor cocktails (Roche) and lysed overnight in -80 °C. Lysates were clarified by centrifugation at 20,000 g for 20 min at 4 °C the next day, and protein concentrations were measured using the BCA protein assay (Pierce). Equal amounts of protein were loaded onto 4-20% precast polyacrylamide gels (Bio-Rad), separated by electrophoresis, and transferred to nitrocellulose membranes (Bio-Rad). Membranes were blocked for 1 h at room temperature in Intercept (TBS) Blocking Buffer (LI-COR) and probed overnight at 4 °C with primary antibodies targeting Wee1 (Cell Signaling Technology, #13084S), CK1α (Abcam, #ab108296), eRF3/GSPT1 (Abcam, #ab49878), Ikaros (Cell Signaling Technology, #14859S), Helios (Cell Signaling Technology, #42427S), Aiolos (Cell Signaling Technology, #15103), p53 (Cell Signaling Technology, #9282), p21 (Cell Signaling Technology, #2947), CRBN (Novus Biologicals, #NBP1-91810), α-Tubulin (Cell Signaling Technology, #3873S), and β-Actin (Cell Signaling Technology, #3700). Following primary antibody incubation, membranes were incubated with IRDye800-labeled goat anti-rabbit IgG or IRDye680-labeled goat anti-mouse IgG secondary antibodies (LI-COR) for 1 h at room temperature. Bands were visualized using the Odyssey CLx imaging system (LI-COR).

### Global label free quantitative proteomics

#### Sample preparation

Cells were lysed by addition of lysis buffer (8 M Urea, 50 mM NaCl, 50 mM 4-(2-hydroxyethyl)-1- piperazineethanesulfonic acid (EPPS) pH 8.5, Protease and Phosphatase inhibitors) and homogenization by bead beating (BioSpec) for three repeats of 30 sec at 2400 strokes/min. Bradford assay was used to determine the final protein concentration in the clarified cell lysate. Fifty micrograms of protein for each sample was reduced, alkylated and precipitated using methanol/chloroform as previously described^23^ and the resulting washed precipitated protein was allowed to air dry. Precipitated protein was resuspended in 4 M urea, 50 mM HEPES pH 7.4, followed by dilution to 1 M urea with the addition of 200 mM EPPS, pH 8. Proteins were digested with the addition of LysC (1:50; enzyme:protein) and trypsin (1:50; enzyme:protein) for 12 h at 37 °C. Sample digests were acidified with formic acid to a pH of 2-3 before desalting using C18 solid phase extraction plates (SOLA, Thermo Fisher Scientific). Desalted peptides were dried in a vacuum-centrifuged and reconstituted in 0.1% formic acid for liquid chromatography-mass spectrometry analysis.

Data were collected using a TimsTOF Pro2 or timsTOF HT (Bruker Daltonics, Bremen, Germany) coupled to a nanoElute LC pump (Bruker Daltonics, Bremen, Germany) via a CaptiveSpray nano-electrospray source. Peptides were separated on a reversed-phase C_18_ column (25 cm x 75 µm ID, 1.6 µM, IonOpticks, Australia) containing an integrated captive spray emitter. Peptides were separated using a 50 min gradient of 2 - 30% buffer B (acetonitrile in 0.1% formic acid) with a flow rate of 250 nL/min and column temperature maintained at 50 °C. The TIMS elution voltages were calibrated linearly with three points (Agilent ESI-L Tuning Mix Ions; 622, 922, 1,222 m/z) to determine the reduced ion mobility coefficients (1/K_0_).

To perform diaPASEF on the TimsTOF Pro2, the precursor distribution in the DDA m/z-ion mobility plane was used to design an acquisition scheme for Data-independent acquisition (DIA) data collection which included two windows in each 50 ms diaPASEF scan. Data was acquired using sixteen of these 25 Da precursor double window scans (creating 32 windows) which covered the diagonal scan line for doubly and triply charged precursors, with singly charged precursors able to be excluded by their position in the m/z-ion mobility plane. These precursor isolation windows were defined between 400 - 1200 m/z and 1/k0 of 0.7 - 1.3 V.s/cm^2^.

To perform diaPASEF on the timsTOF HT, we used py_diAID (Skowronek et al., 2022), a python package, to assess the precursor distribution in the m/z-ion mobility plane to generate a diaPASEF acquisition scheme with variable window isolation widths that are aligned to the precursor density in m/z. Data was acquired using twenty cycles with three mobility window scans each (creating 60 windows) covering the diagonal scan line for doubly and triply charged precursors, with singly charged precursors able to be excluded by their position in the m/z-ion mobility plane. These precursor isolation windows were defined between 350 - 1250 m/z and 1/k_0_ of 0.6 - 1.45 V.s/cm^2^.

#### LC-MS data analysis

The diaPASEF raw file processing and controlling peptide and protein level false discovery rates, assembling proteins from peptides, and protein quantification from peptides were performed using either a targeted cell line specific spectral library generated by searching offline fractionated DDApasef data against a Swissprot human database (January 2021) or library free analysis in DIA-NN 1.8.^24^ Library free mode performs an *in silico* digestion of a given protein sequence database alongside deep learning-based predictions to extract the DIA precursor data into a collection of MS2 spectra. The search results are then used to generate a spectral library which is then employed for the targeted analysis of the DIA data searched against a Swissprot human database (January 2021). Database search criteria largely followed the default settings for directDIA including: tryptic with two missed cleavages, carbamidomethylation of cysteine, and oxidation of methionine and precursor Q-value (FDR) cut-off of 0.01. Precursor quantification strategy was set to Robust LC (high accuracy) with RT-dependent cross run normalization. Proteins with low sum of abundance (<2,000 x no. of treatments) were excluded from further analysis and resulting data was filtered to only include proteins that had a minimum of 3 counts in at least 4 replicates of each independent comparison of treatment sample to the DMSO control. Protein with missing values were imputed by random selection from a Gaussian distribution either with a mean of the non-missing values for that treatment group or with a mean equal to the median of the background (in cases when all values for a treatment group are missing) using in-house scripts in the R framework (R Development Core Team, 2014). Significant changes comparing the relative protein abundance of these treatment to DMSO control comparisons were assessed by moderated t test as implemented in the limma package within the R framework.^25^

### Molecular docking

Docking studies were carried out with the Maestro program from Schrödinger. CK1α + CRBN + DDB1 + **SJ3149** quaternary complex (PDB Code: 8G66) was used as the protein template.^17^ Protein Preparation and LigPrep wizards were applied to prepare the protein and ligands by using the default parameters. Docking grid was generated with the Receptor Grid Generation module. Hydrogen bond constraints between the ligand and the backbone carbonyl oxygen and imidazole NH of CRBN His378, backbone NH of CRBN Trp380 were set as the constraints. Then Ligand Docking was conducted with standard precision. All the three hydrogen bond constraints must be matched during the docking. Default values were adopted for other parameters.

### Chemical synthesis

Additional details are provided in the supporting information

## Acknowledgements

This work was supported by the National Institutes of Health (NIH) grant NIH High End Instrumentation grant (1S10OD028697-01) and R01CA214608 (to N.S.G and E.S.F.), NIH career development award (K08CA23022) (to S.M.C.), and departmental funds from Stanford Chemical and Systems Biology and Stanford Cancer Institute (to N.S.G.).

W.S.B. is supported by Basic Science Research Program through the National Research Foundation of Korea (NRF) funded by the Ministry of Education (grant no. RS-2024-00410290).

R.C.S. acknowledges the Swiss National Science Foundation for a postdoctoral fellowship (SNF Mobility grant P500PN_206898).

---

**Figure S1.**
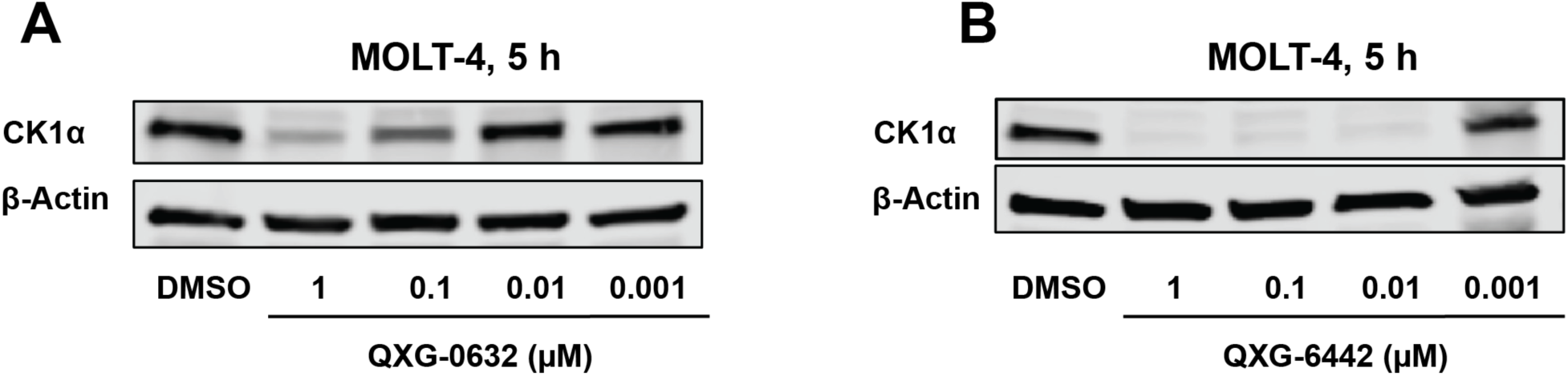
Western blot analysis of CK1α treated with the indicated concentration of **QXG-0632** or **QXG-6442** for 5 h.

**Figure S2.**
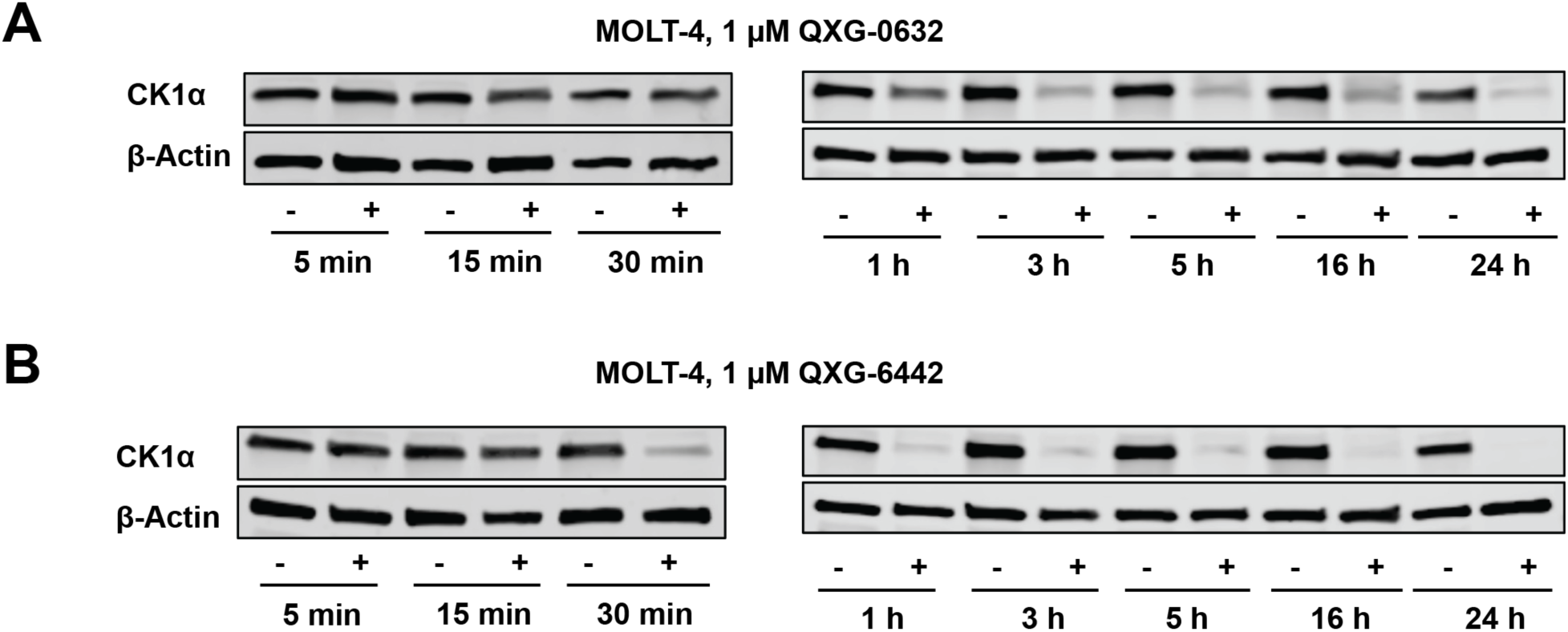
Western blot analysis of CK1α treated with 1 μM of **QXG-0632** or **QXG-6442** for the indicated time.

**Figure S3.**
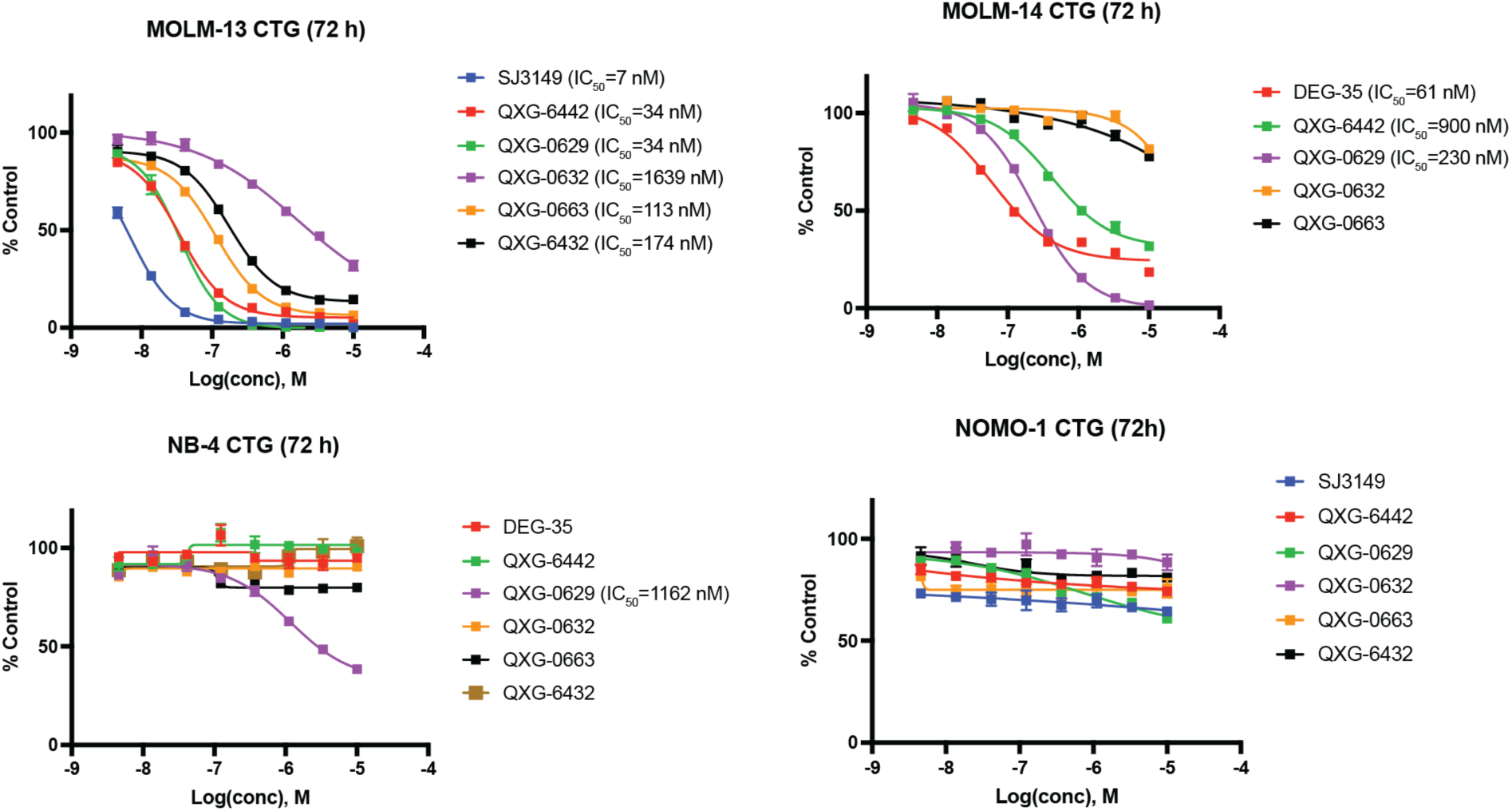
Antiproliferative effect of potent CK1α degraders in representative types of cancer cell lines.

**Figure S4.**
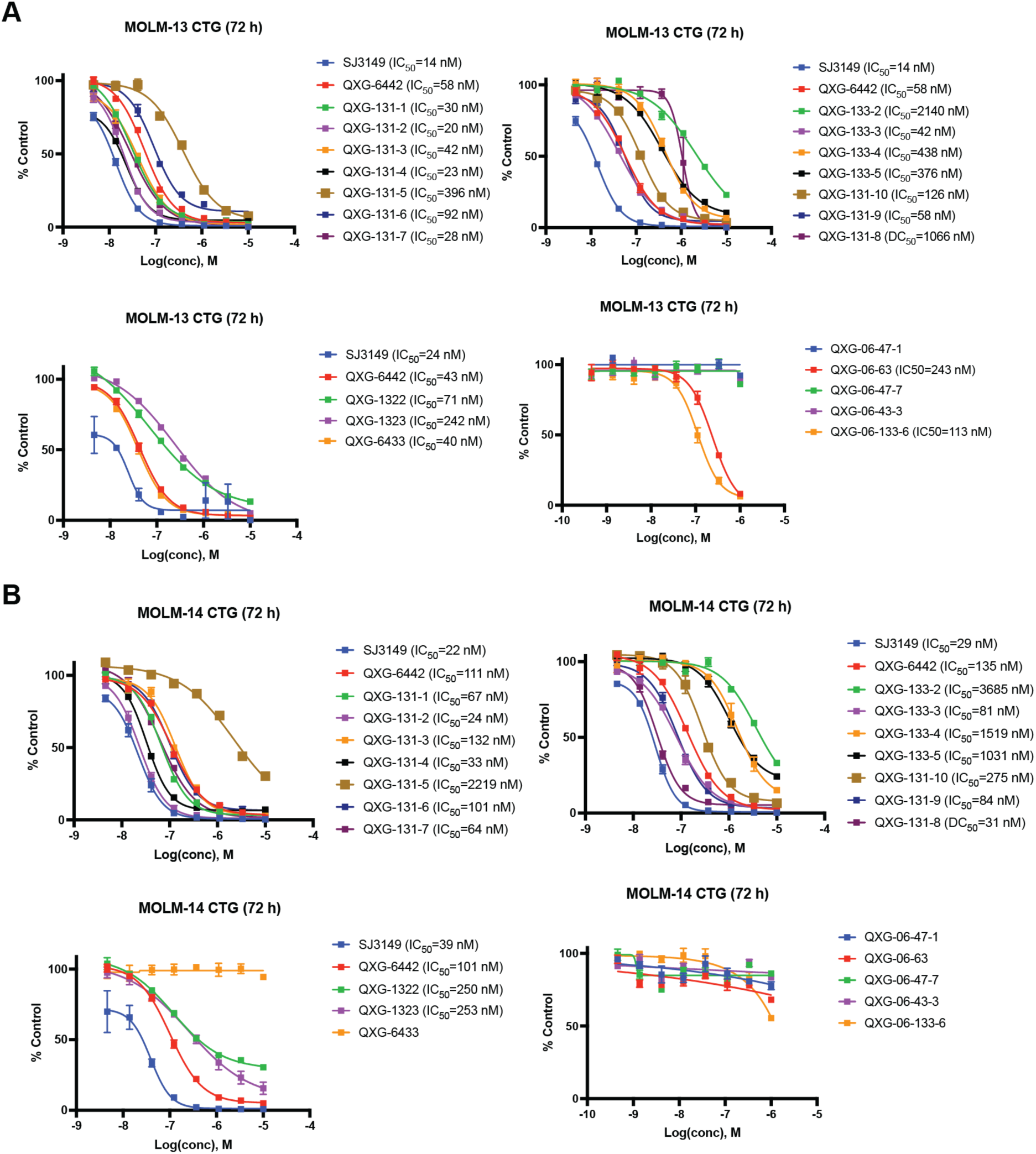
(A) Antiproliferative effect of **QXG-6442** analogs in MOLM-13. (B) Antiproliferative effect of **QXG-6442** analogs in MOLM-14.

**Figure S5.**
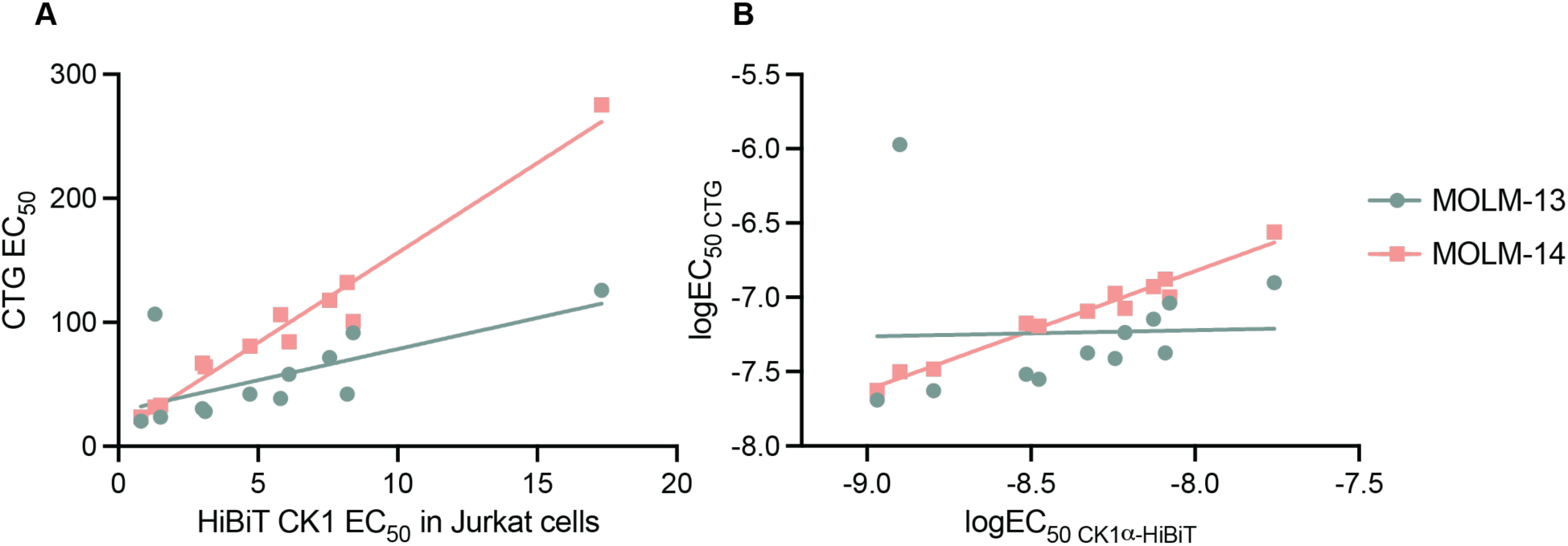
Correlation of cytotoxicity (EC_50_, CTG in A; logEC_50_, CTG in B) and degradation activity (logDC_50_, HiBiT in A; logDC_50_, HiBiT in B) results modeled with a linear regression (MOLM-13 R squared = 0.0011, MOLM-14 R squared = 0.97).

**Figure S6.**
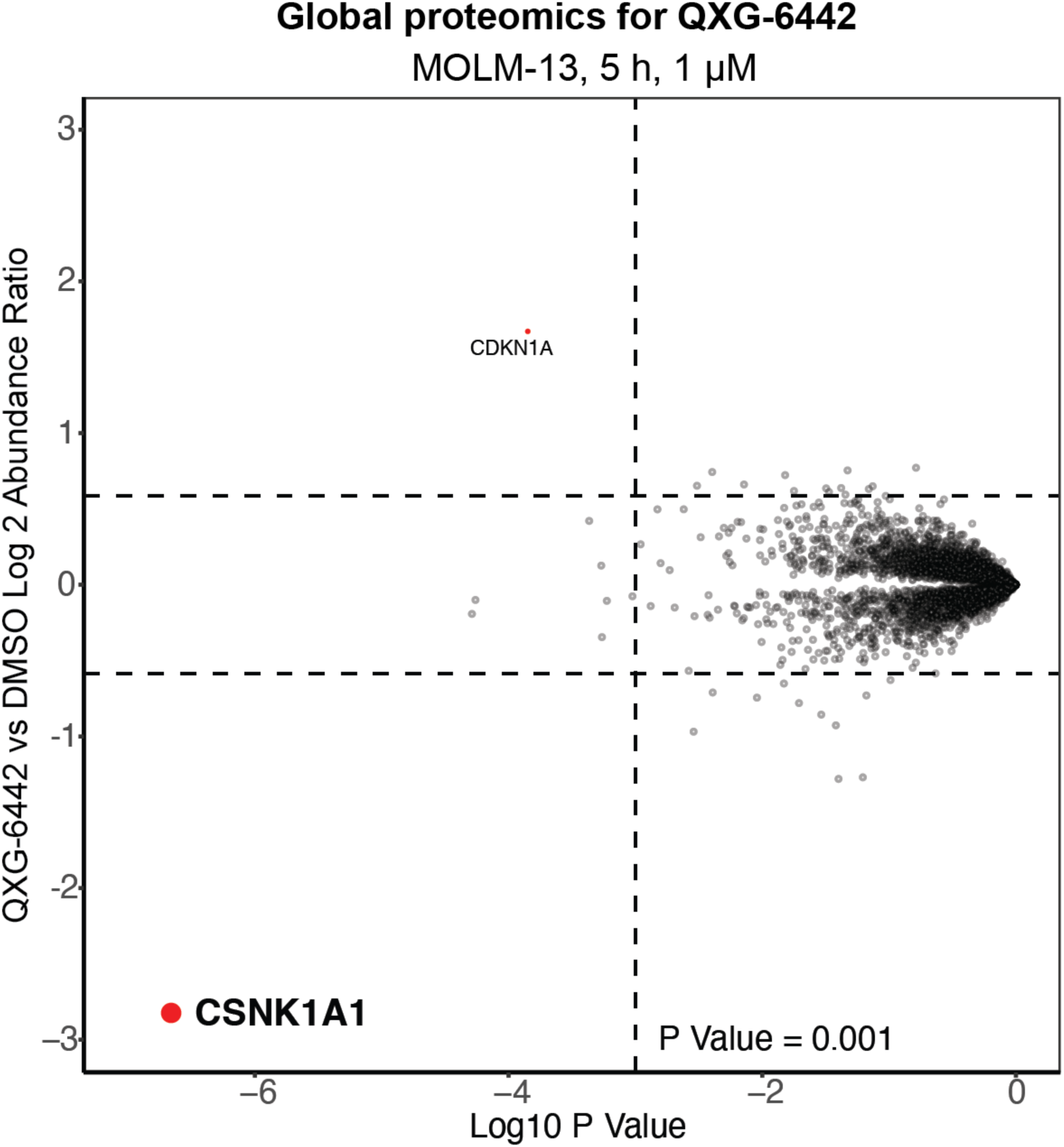
Global proteomics analysis of **QXG-6442** in MOLM-13.

**Table S1.**
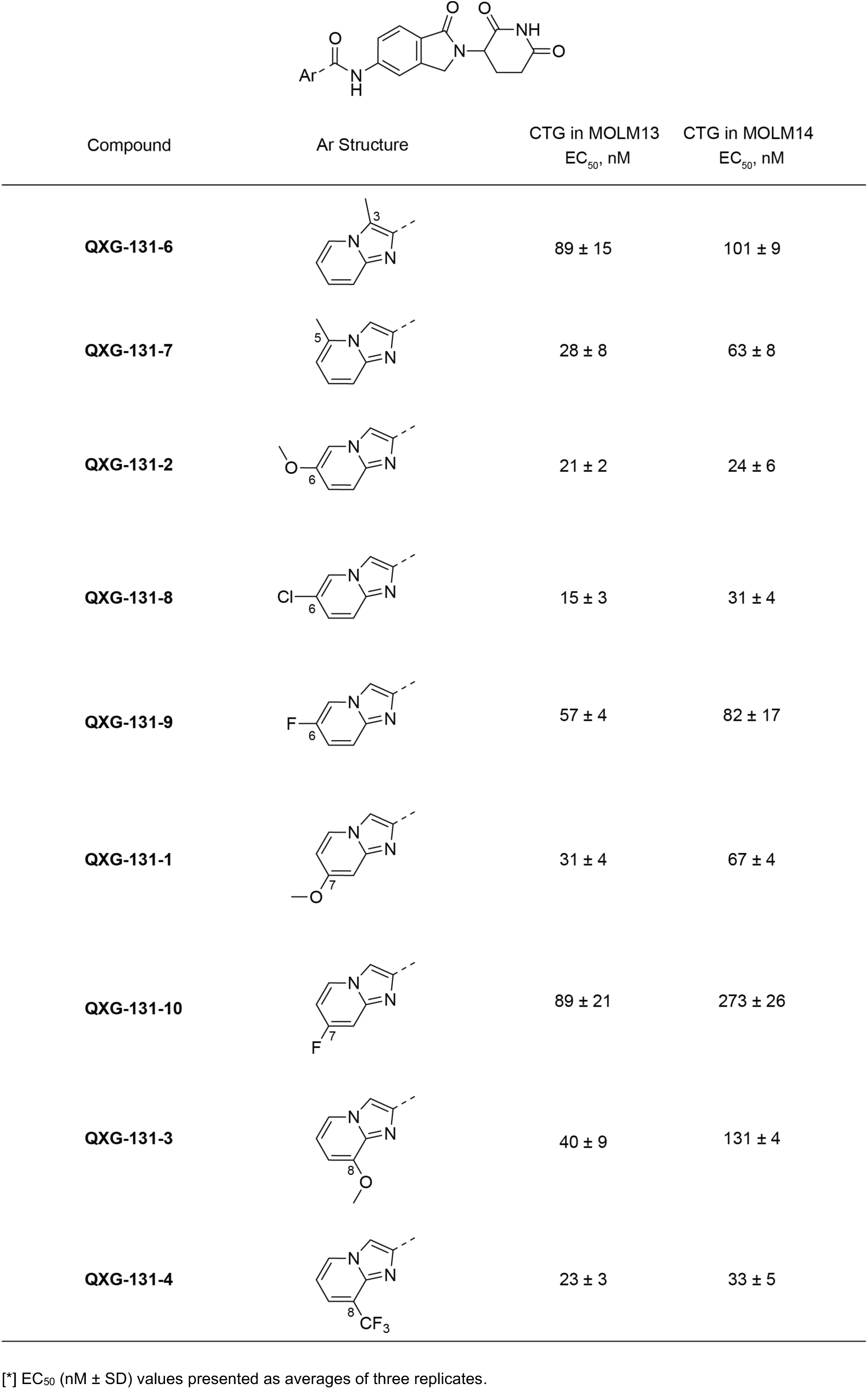
Structure-activity relationship of antiproliferative effects of **QXG-6442** analogs

**Table S2.**
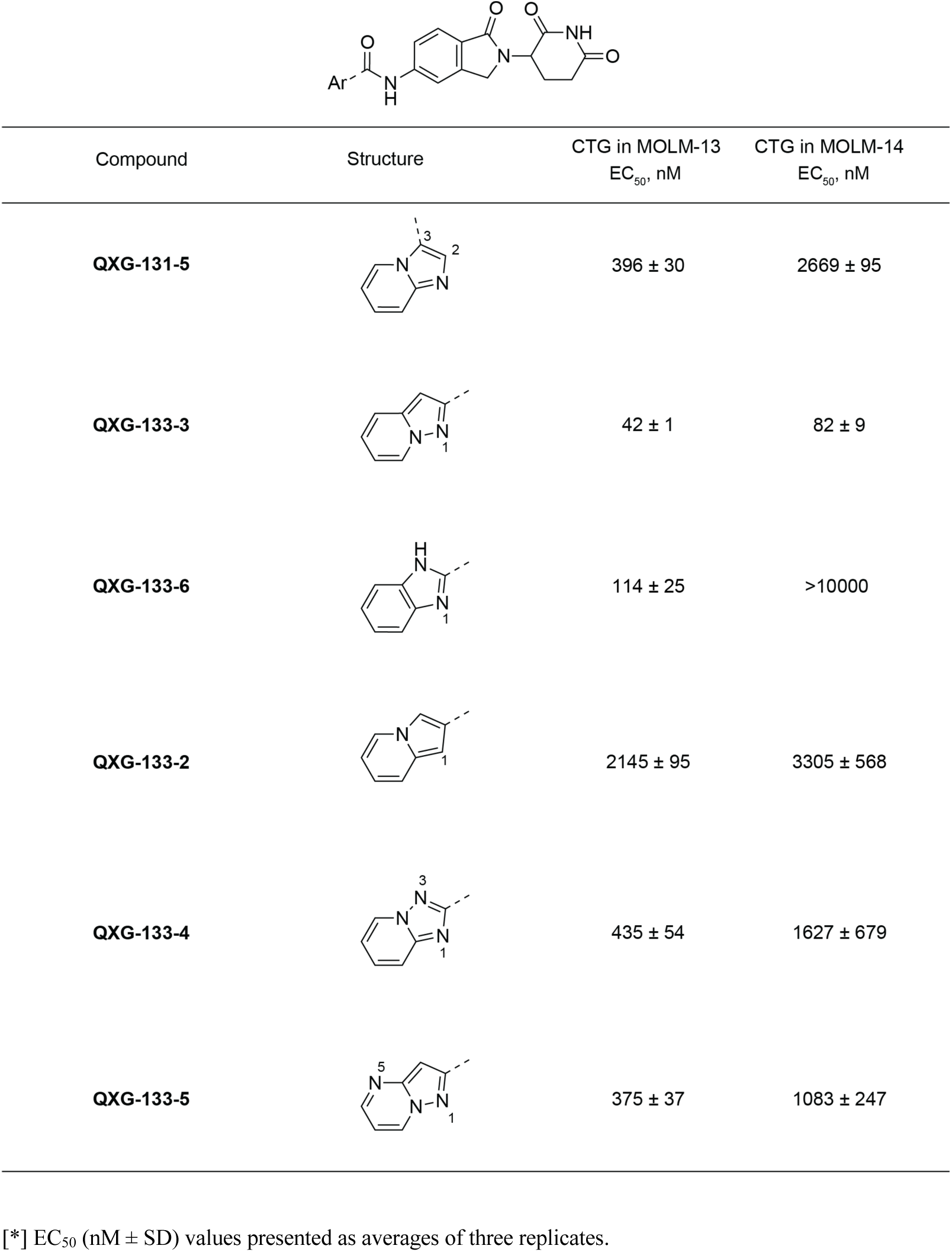
Structure-activity relationship of antiproliferative effects for different gluing moiety derivated from **QXG-6442**

| Compound | Structure | CTG in MOLM-13<br>EC<sub>50</sub>, nM | CTG in MOLM-14<br>EC<sub>50</sub>, nM |
| --- | --- | --- | --- |
| **QXG-131-5** |  | 396 ± 30 | 2669 ± 95 |
| **QXG-133-3** |  | 42 ± 1 | 82 ± 9 |
| **QXG-133-6** |  | 114 ± 25 | >10000 |
| **QXG-133-2** |  | 2145 ± 95 | 3305 ± 568 |
| **QXG-133-4** |  | 435 ± 54 | 1627 ± 679 |
| **QXG-133-5** |  | 375 ± 37 | 1083 ± 247 |

[\*] EC<sub>50</sub> (nM ± SD) values presented as averages of three replicates.

